# *In vitro* cultivation techniques for modeling liver organogenesis, building assembloids, and designing synthetic tissues

**DOI:** 10.1101/2023.09.30.560154

**Authors:** Simran Kumar, Helly Patel, Natesh Parashurama

## Abstract

Chronic liver disease has reached epidemic proportions, affecting over 800 million people globally. The current treatment, orthotopic liver transplantation, has several limitations. Promising solutions have emerged in the field of liver regenerative medicine, with liver organogenesis holding significant potential. Early liver organogenesis, occurring between E8.5 and 11.5, involves the formation of epithelial-mesenchymal interactions leading to morphogenesis, hepatic cord formation, and collective migration. However, there is a lack of methods for *in vitro* modeling of this process. In this study, a detailed series of methods are presented enabling the modeling of various stages and aspects of liver organogenesis. In one method series, assembloid technology with hepatic and mesenchymal spheroids is utilized, replicating early structures found in liver organogenesis, modeling early morphogenesis, and demonstrating interstitial cell migration as seen *in vivo*. These innovative assembloid systems help identify factors influencing assembloid formation and migration. Hepatic spheroid cultivation systems were also employed to model collective migration and branching morphogenesis. Fibroblast-conditioned media (MES-CM) plays a significant role in initiating dose-dependent branching migration. Future work will involve high temporal and spatial resolution imaging of hepatic and mesenchymal interactions to determine the cascade of cellular and molecular events involved in tissue formation, morphogenesis, and migration.

**SUMMARY:** Organoids revolutionize personalized tissue modeling for organ development, drug discovery, and disease research. Organoid engineering advances into creating intricate synthetic tissues. The aim is to integrate morphogenesis, assembloid technology, and biomatrices to advance tissue engineering. The presented methods aid in modeling liver organogenesis and establishing guidelines for synthetic tissue construction.

## INTRODUCTION

Liver cell migration plays a significant role in liver organogenesis, disease, and cell therapy. During liver organogenesis (E8.5-9.0, mouse), the ventral foregut pre-hepatic epithelium begins to express liver genes due to inductive signals emanating from the surrounding mesenchyme and heart. At E9.0, the foregut epithelium thickens as the cells transition from cuboidal to pseudostratified columnar morphology, forming the liver diverticulum ^3, 7^. At this critical stage, the liver diverticulum is comprised of only ∼1,500 cells. Next, the hepatic endoderm lining the liver diverticulum thickens, delaminates, and forms cords of migrating hepatoblasts that co-migrate with endothelial cells and mesenchymal cells, branching into the surrounding mesenchymal tissue, initiating three-dimensional collective cell migration to form the liver bud ^15, 16^. In fact, during this stage, cells collectively undergo: 1) co-migration, or movement together with other cell types, 2) branching morphogenesis, or formation of branching tube-like structures, and 3) interstitial migration, or migration on top of other cells. By E11.5, migration ceases, and the primitive liver has formed, expanding 10^3^-fold ^15^. Liver cell migration may also be required in later stages of liver organogenesis, as rat fetal hepatoblasts (HBs) expression has shown evidence of highly upregulated genes associated with 3D collective cell migration, morphogenesis, and extracellular matrix remodeling^18^. In addition to its role in early liver organogenesis, 3D collective migration is intricately linked to the local spread and metastasis of advanced hepatocellular carcinoma (HCC), ultimately leading to worsened prognosis and increased treatment resistance^25^. Adult and fetal hepatocytes also employ collective migration when moving from the spleen to within the liver during liver repopulation. *In vivo* imaging studies have demonstrated that transplanted hepatocytes enter the portal vein and then the capillaries within hours, migrate across the liver sinusoids, and through the liver tissue^8, 10, 19^. Finally, recent studies demonstrate that migrating hepatoblasts arise during murine and human liver regeneration with some evidence of movement in sheets^11^. Overall, liver collective migration, capable of multiple modes of morphogenesis, plays a significant role in organogenesis, cancer, hepatocyte cell therapy, and liver regeneration.

Numerous genetic studies have investigated the molecular pathways that drive 3D liver collective cell migration. These studies demonstrate that ablation of hepatic cords blocks liver formation and shows that, therefore, the formation of hepatic cords and their ensuing interactions with supporting cells are required for liver formation^3, 12, 21, 22^. These studies also demonstrate that liver growth is initiated by fibroblast growth factor 2 (FGF2) secreted from the cardiac mesoderm, bone morphogenetic protein 4 (BMP4) secreted from the surrounding mesenchyme, hepatocyte growth factor (HGF), endothelial cell interactions, and migration-associated transcription factors including (HEX), PROX1, and TBX3^7, 20^. Overall, genetic studies support the fact that soluble factor signaling leading to transcription factor expression is responsible for driving migration, signaling, and molecular interactions between hepatoblasts and their surrounding mesenchyme.

Although cell migration in early liver organogenesis has been extensively investigated, current *in vitro* hepatic migration studies frequently utilize 2D assays consisting of highly migratory HCC cells combined with *in vivo* tumor models^13^. These studies have provided insight into several factors that play a role in hepatic migration including TGFβ1^6^, c-Myc^27^, Yes-associate protein (YAP)^5^, goosecoid^23^, actopaxin^2^, and miRNAs^4, 24, 26^. Despite the advancements in understanding the molecular mechanisms in 3D hepatic cell migration, the fundamental mechanisms between 2D and 3D cellular migration are distinct, suggesting 2D assays have their limitations. Furthermore, these models typically do not implement mesenchymal cell types, which are essential to migration/growth. There has been progress in the development of 3D models for liver migration that incorporate the supporting mesenchyme; however, they are solely focused on co-migration rather than the different modes of collective migration.

The ability to form tissues from spheroids through various self-assembly and morphogenetic processes enables the scientific study of synthetic tissues for applications in drug development and screening, disease modeling, therapy, and other biomedical and biotechnological applications. Methodological details for several 3D *in vitro* cultivation systems engineered to exhibit different modes of liver 3D collective migration are presented. These systems include the following: (1) co-spheroid culture with hepatic (HEP) and mesenchymal-derived (MES) spheroids in matrix, (2) spheroid matrix droplet cultured with fibroblast-conditioned medium (MES-CM), and (3) mixed spheroids (HEP and MES cells). These systems enable robust modeling of liver 3D collective migration, improving our molecular and cellular understanding of liver organogenesis, cancer, and therapy.

## PROTOCOL

### 1. Preparation of 1% Low EEO Agarose Solution

1.1. Weigh 2.5 g of low EEO agarose powder and transfer it to a beaker.

NOTE: Ensure the beaker is at least twice the size of the desired volume to accommodate bubbling during the solution preparation.

1.2. Use a graduated cylinder to measure 250 mL of distilled (DI) water and pour it into the beaker, aiming for a final concentration (w/v) of 1%.

1.3. Cover the beaker’s opening with plastic wrap, leaving a small hole for ventilation. Heat the beaker in the microwave for 30 seconds.

1.4. Carefully remove the beaker and swirl the contents. Repeat this process every 30 seconds until complete dissolution.

CAUTION: Handle microwaved glassware carefully, using proper gloves. Monitor the solution closely to prevent overheating or boiling over.

1.5. Once a homogeneous solution is achieved, remove the beaker from the microwave. Gently swirl the solution before transferring it to a pre-sterilized bottle. Autoclave the solution and store the agarose solution at room temperature until ready to use.

### 2. Coating 96-Well Plate

NOTE: 384-well ultra-low attached spheroid microplates can be utilized in these and further experiments.

2.1. Loosen the cap of the bottle containing the sterile 1% agarose solution. Warm the solution in the microwave in 30-second intervals until it transitions to the liquid phase; then tighten the cap.

CAUTION: Handle microwaved glassware carefully, using proper gloves. Monitor the solution closely to prevent overheating or boiling over.

NOTE: Perform these steps under a sterile tissue-culture laminar flow hood.

2.2. Transfer 55-65 µL of the 1% agarose solution per well to coat the 96-well tissue-cultured plate and immediately rotate the plate.

2.3. Once the 1% agarose solution has been transferred to the desired number of wells, place the plates in a 4℃ fridge for 20-30 minutes to allow the agarose to solidify. Before use, bring the plate to room temperature (**Figure 1A**).

**Figure 1.**
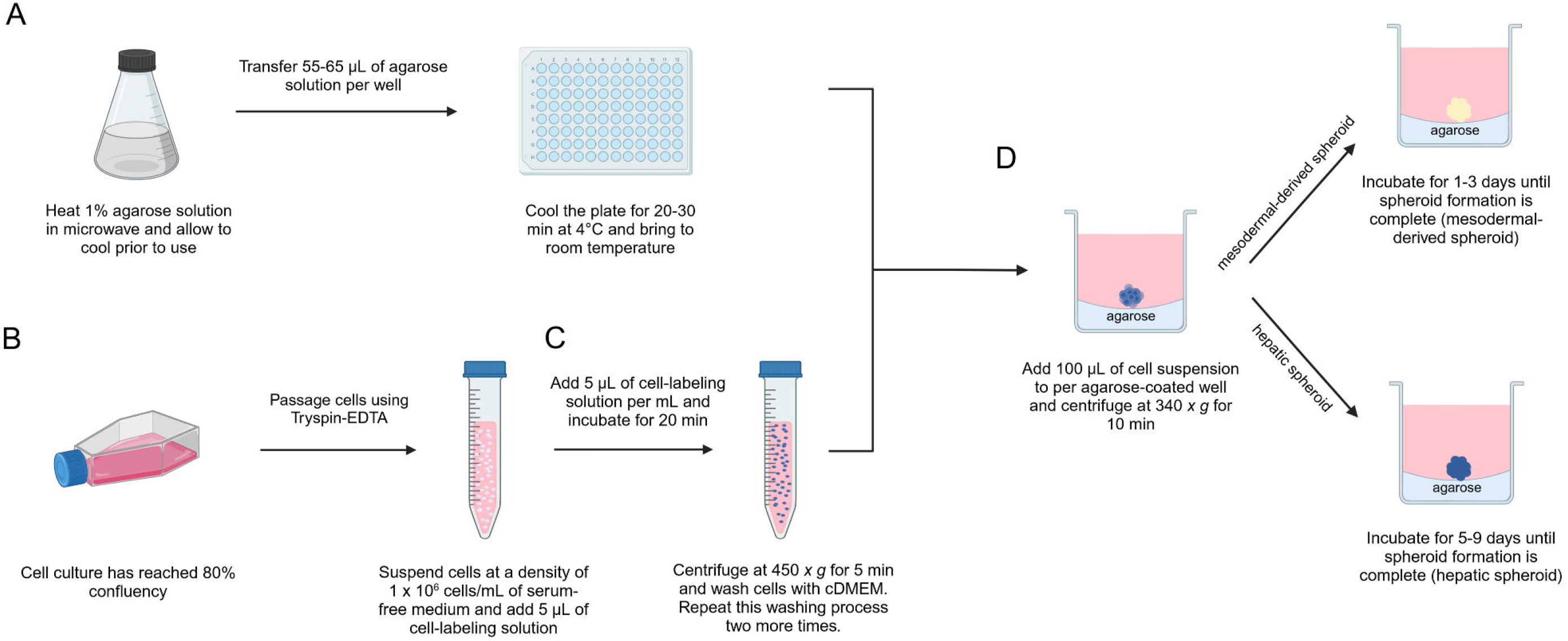
Schematic of spheroid formation assay. **(A)** Coating a 96-well plate with 1% agarose solution. Agarose solution is melted in the microwave, 55-65 µL is transferred to each well and the plate is cooled for 20-30 minutes. **(B)** Passaging of cell culture at 80% confluency. **(C)** Dye-labeling of cell suspension. 5 µL of cell-labeling solution is added per mL After incubation and centrifugation, the cell suspension is washed with warm cDMEM before plating the cells. **(D)** Plating of cell suspension for spheroid formation. 100 µL of cell suspension is added to each well and the plate is centrifuged. Mesodermal-derived spheroids form in 1-3 days, while hepatic spheroids form in 5-9 days.

### 3. Preparation of HepG2 (HEP) Spheroids

3.1. Cultivate HepG2 cells in a T-75 flask using completed growth medium (cDMEM) comprising of Dulbecco’s Modified Eagle Medium (DMEM), supplemented with 10% Fetal Bovine Serum (FBS) and 1% Penicillin-Streptomycin (Pen-Strep). Maintain the cell culture at 37℃ and 5% CO_2_, changing the medium daily.

3.2. Upon reaching 80% confluency, perform a rinse of the cells with 1X Phosphate Buffered Saline (PBS) and discard the rinse. Add 5 mL of 0.05% of Trypsin-EDTA to the flask and incubate at 37℃ and 5% CO_2_ for 5-10 minutes.

3.3. Add equal amounts of cDMEM and wash the cells off the flask. Once cells detach, transfer the mixture to a 15 mL sterile conical centrifuge tube and centrifuge the cell suspension at 300 *x g* for 5 minutes.

NOTE: All centrifugation steps outlined in this protocol were performed at room temperature. However, these centrifugation procedures can alternatively be carried out at cold temperatures.

3.4. Re-suspend the cell suspension in fresh cDMEM and count the cells to determine the final concentration in cells/mL (**Figure 1B**).

3.5. Dye-labeling of cells

NOTE: This is an optional step.

3.5.1. Transfer the desired amount of cell suspension to a 15 mL sterile conical centrifuge tube and centrifuge the cell suspension at 300 *x g* for 5 minutes.

3.5.2. Re-suspend the cell pellet in a serum-free growth medium to achieve a final concentration of 1.0 x 10^6^ cells/mL. Add 5 µL of Vybrant Cell-Labeling Solution per mL of cell suspension and incubate the cell suspension at 37℃ and 5% CO_2_ on rotation for 20 minutes.

NOTE: Serum-free growth medium is DMEM only supplemented with 1% Pen-strep. Adjust the incubation time for uniform staining based on the cell suspension density.

3.5.3. Once the incubation is completed, centrifuge the cell suspension at 450 *x g* for 5 minutes and re-suspend the cell pellet in fresh cDMEM. Repeat this wash process two more times (**Figure 1C**).

3.6. Spheroid formation

3.6.1. Suspend the cells in fresh cDMEM to achieve a final concentration of 5.0 x 10^4^ cells/mL based on the cell count. Mix the cell suspension well and transfer 100 µL of cell suspension per well to the agarose-coated 96-well plate.

NOTE: The cell suspension density is set to achieve 5,000 cells per well in the agarose-coated 96-well plate.

3.6.2. Centrifuge the plate at 340 *x g* for 10 minutes and incubate at 37℃ and 5% CO2 for 5-9 days (**Figure 1D**).

NOTE: Change medium every other day after plating, gently removing 50% of cDMEM and replacing it.

NOTE: HepG2 spheroids are suitable for the HepG2 and HFF/MRC-5 (HEP-MES) assembloid model or MES-CM model.

### 4. Preparation of HFF/MRC-5 (MES) Spheroids

4.1. Cultivate HFF/MRC-5 cells in a T-175 flask using completed growth medium (cDMEM) comprising of Dulbecco’s Modified Eagle Medium (DMEM), supplemented with 10% Fetal Bovine Serum (FBS) and 1% Penicillin-Streptomycin (Pen-Strep). Maintain the cell culture at 37℃ and 5% CO_2_, changing the medium every other day.

4.2. Upon reaching 80% confluency, perform a rinse of the cells with 1X Phosphate Buffered Saline (PBS) and discard the rinse. Add 5 mL of 0.25% of Trypsin-EDTA to the flask and incubate at 37℃ and 5% CO2 for 5-10 minutes.

4.3. Add equal amounts of cDMEM and wash the cells off the flask. Once cells detach, transfer the mixture to a 15 mL sterile conical centrifuge tube and centrifuge the cell suspension at *300 x g* for 5 minutes.

4.4. Re-suspend the cell suspension in fresh cDMEM and count the cells to determine the final concentration in cells/mL (**Figure 1B**).

4.5. Dye-labeling of Cells

NOTE: This is an optional step.

4.5.1. Transfer the desired amount of cell suspension to a 15 mL sterile conical centrifuge tube and centrifuge the cell suspension at 300 *x g* for 5 minutes.

4.5.2. Re-suspend the cell pellet in a serum-free growth medium to achieve a final concentration of 1.0 x 10^6^ cells/mL. Add 5 µL of Vybrant Cell-Labeling Solution per mL of cell suspension and incubate the cell suspension at 37℃ and 5% CO2 on rotation for 20 minutes.

NOTE: Serum-free growth medium is DMEM only supplemented with 1% Pen-strep. Adjust the incubation time for uniform staining based on the cell suspension density.

4.5.3. Once the incubation is completed, centrifuge the cell suspension at 450 *x g* for 5 minutes and re-suspend the cell pellet in fresh cDMEM. Repeat this wash process two more times (**Figure 1C**).

4.6. Spheroid Formation

4.6.1. Suspend the cells in fresh cDMEM to achieve a final concentration of 1.0 x 10^5^ cells/mL based on the cell count. Mix the cell suspension well and transfer 100 µL of cell suspension per well to the agarose-coated 96-well plate.

NOTE: The cell suspension density is set to achieve a density of 10,000 cells per well in the agarose-coated 96-well plate.

4.6.2. Centrifuge the plate at 340 *x g* for 10 minutes and incubate at 37℃ and 5% CO2 for 5-9 days (**Figure 1D**).

NOTE: Change medium every other day after plating, gently removing 50% of cDMEM and replacing it.

NOTE: HFF/MRC-5 spheroids are suitable for the HEP-MES assembloid model.

### 5. HepG2 and HFF/MRC-5 (HEP-MES) Assembloid Formation

NOTE: Refer to Sections 3 and 4 for formation of HepG2 (**Figure 2A)** and HFF/MRC-5 spheroids (**Figure 2B**).

**Figure 2.**
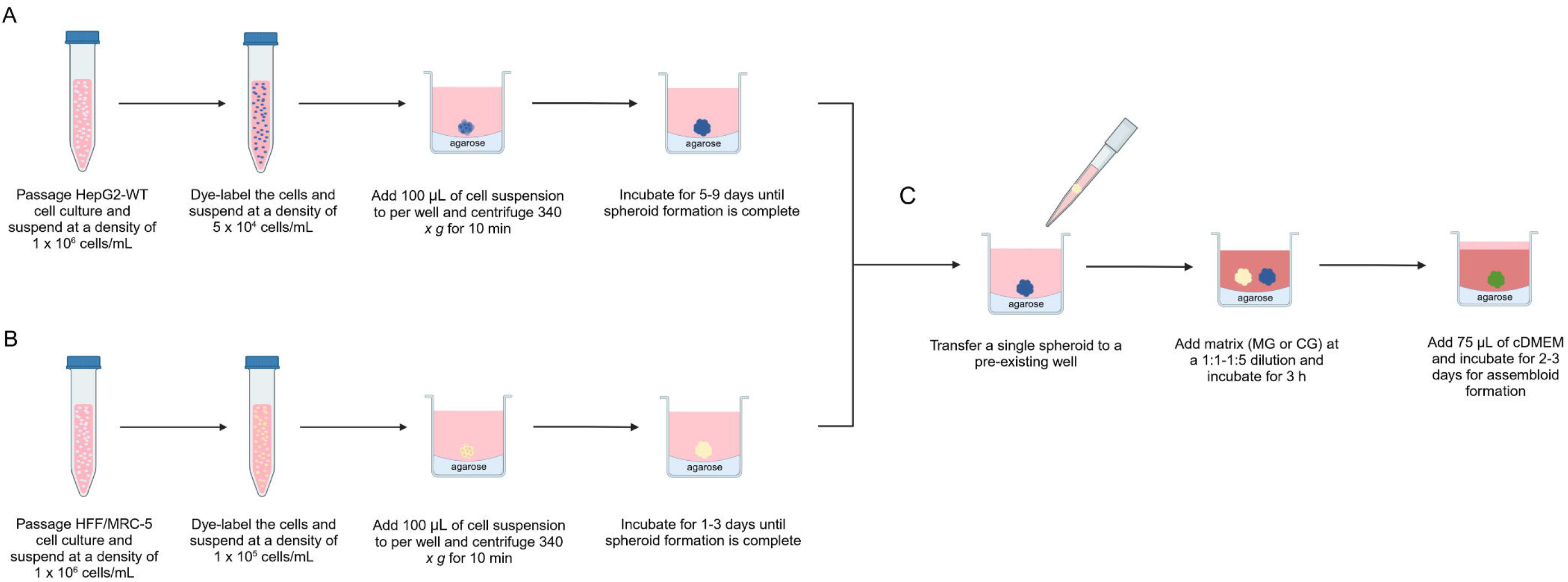
Schematic of assembloid formation assay with hepatic and mesodermal-derived spheroids. **(A)** Hepatic spheroid formation. HepG2 cells are dye-labeled and plated for spheroid formation. **(B)** Mesodermal-derived spheroid formation. HFF/MRC-5 cells are dye-labeled and plated for spheroid formation. **(C)** HEP-MES assembloid formation. Transfer mesodermal-derived spheroid to hepatic spheroid for assembloid formation. Matrix (MG or CG) is added (1:1-1:5 dilution), incubated for 3 hours, followed by the addition of 75 μL cDMEM. Incubated for 2-3 days until assembloid formation is complete.

5.1. Collect HFF/MRC-5 spheroids individually using a pipette from the 96-well plate. Transfer them to a 15 mL sterile conical centrifuge tube containing fresh cDMEM. Allow the spheroids to settle in the tube for 5 minutes and then gently rinse with warm cDMEM.

NOTE: Perform the rinsing process gently and with care.

5.2. Gently transfer a single HFF/MRC-5 spheroid using a pipette to a well containing a HepG2 spheroid. Add ice-cold growth factor reduced (GFR) Matrigel (MG)/ Collagen (CG) in a 1:1 and 1:5 dilution and incubate at 37°C at 5% CO_2_ for 3 hours (**Figure 2C**).

NOTE: Thaw MG overnight on ice in a 2°C to 8°C fridge. Ensure the use of pre-cooled pipettes, tips, and tubes when handling MG, as it forms a gel above 10℃.

5.3. After incubation, add 75 µL of fresh cDMEM to each well and incubate at 37°C at 5% CO_2_ for 2-3 days.

NOTE: Change medium every other day after plating, gently removing 50% of cDMEM and replacing it. Assembloids will still form without media changes for up to 3-4 days.

NOTE: Assembloid formation can occur without the use of matrix.

### 6. HEP Spheroid Droplet Formation

NOTE: Refer to Section 3 for formation of HepG2 spheroids (**Figure 3A**).

**Figure 3.**
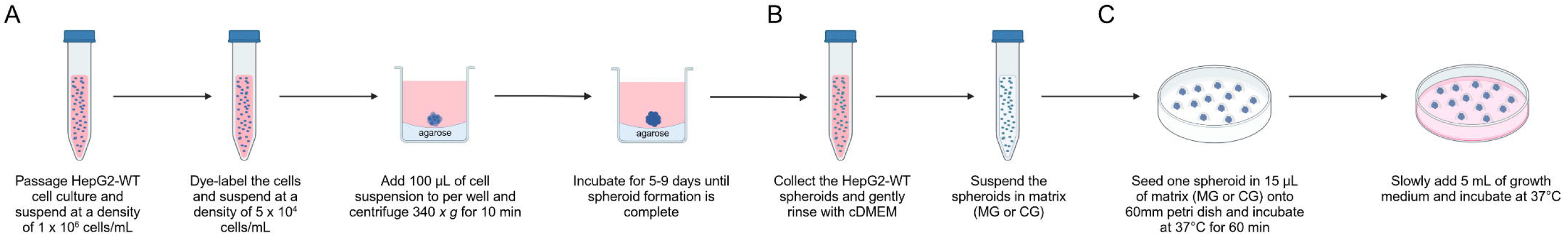
Schematic of Droplet Formation Assay. **(A)** Hepatic spheroid formation. HepG2 cells are dye-labeled. **(B)** Spheroid collection and matrix suspension. Collect and gently rinse hepatic spheroids with cDMEM. Spheroids are suspended in MG or CG solution (1:1 – 1:5 dilution). **(C)** Droplet seeding and addition of growth medium. Hepatic spheroids are seeded onto a 60mm petri dish and incubated for 60 minutes, and 5 mL of MES-CM is added. Petri dish is incubated with media changes every three days.

NOTE: Two different materials can be used for suspending the HepG2 spheroids in droplets. The two methods are provided below.

6.1. Matrigel (MG) droplets

6.1.1. Collect HepG2 spheroids individually using a pipette from the 96-well plate, allowing them to settle in a 15 mL sterile conical centrifuge on ice.

6.1.2. In a separate 15 mL sterile conical centrifuge tube, mix 1 mL of ice-cold GFR MG and ice-cold control growth medium at a 1:1 dilution. Aspirate the medium from the 15 mL tube containing the HepG2 spheroids and place it on ice.

NOTE: Thaw MG overnight on ice in a 2°C to 8°C fridge. Ensure pre-cooled pipettes, tips, and tubes when handling MG, as it forms a gel above 10℃.

6.1.3. Add the MG solution to HepG2 spheroids, ensuring even spheroid distribution inside the MG solution.

6.1.4. Using a 200 µL pipette, collect a 15 µL volume of a single spheroid in the MG solution and slowly seed it onto a 60 mm petri dish (**Figure 3B**).

NOTE: If more than one spheroid is seeded per droplet, remove, and reseed properly. Spheroids are repositioned if needed to ensure they are at the center of the droplet.

NOTE: For optimal spheroid seeding, consider using reverse pipetting to minimize air bubble formation.

6.2. Collagen (CG) Droplets

NOTE: All CG preparation should be done on ice.

6.2.1. Collect HepG2 spheroids individually using a pipette from the 96-well plate, allowing them to settle in a 15 mL sterile conical centrifuge tube on ice (**Figure 3B**).

6.2.2. In a microcentrifuge tube, add 358.8 µL of de-ionized water, 100 µL of 10X PBS, 12.1 µL of 1 N NaOH, and 529.1 µL of stock rat tail CG for a total volume of 1 mL.

NOTE: Stock rat tail CG should always be added at the end.

6.2.3. Add the CG solution to HepG2 spheroids, ensuring even spheroid distribution inside the CG solution.

6.2.4. Using a 200 µL pipette, collect a 15 µL volume of a single spheroid in the CG solution and slowly seed it onto a 60 mm petri dish.

NOTE: If more than one spheroid is seeded per droplet, remove, and reseed properly. Spheroids are repositioned to ensure they are at the center of the droplet.

NOTE: For optimal spheroid seeding, consider using reverse pipetting to minimize air bubble formation.

• 6.3 Incubate the droplets at 37℃ and 5% CO2 for 60 minutes before the addition of growth medium.

• 6.4 Slowly add 5 mL of the desired growth medium to the petri dish and incubate at 37°C and 5% CO_2_ with medium changes every three days (**Figure 3C**).

NOTE: HFF/MRC-5 conditioned-media was used in the droplet formation assay.

### 7. Preparation of HFF/MRC-5 Conditioned Media (MES-CM)

7.1. Harvest HFF/MRC-5 cells and seed them into a T-75 tissue culture-treated flask at a seeding density of 5,000 cells/cm^2^ and incubate the flask at 37℃ and 5% CO2 for 72 hours in 15 mL of cDMEM.

NOTE: Monitor the flask during this period to ensure optimal cell health.

7.2. After the 72-hour incubation, collect the fibroblast-conditioned growth medium in a 15 mL sterile conical centrifuge tube. Centrifuge the fibroblast-conditioned growth medium at 290 *x g* for 5 minutes and filter using a 0.2 µm filter (**Figure 4A**).

**Figure 4.**
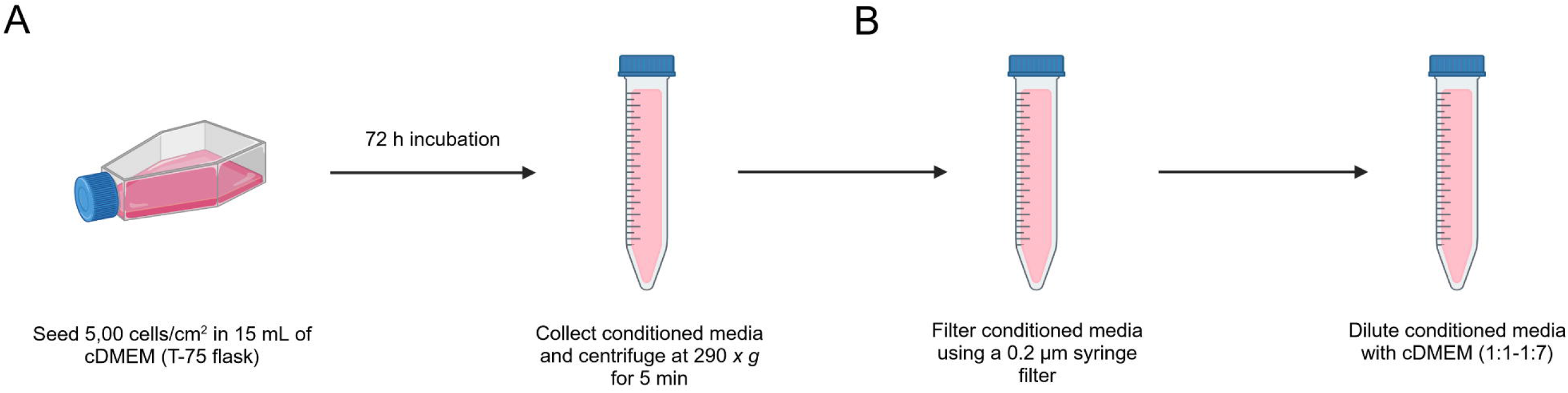
Schematic of Preparation of MES-CM. **(A)** Cell seeding and MES-CM collection. HFF/MRC-5 cells are seeded at 5,000 cells/cm^2^ in a T-75 flask and incubated for 72 hours. The media is collected and centrifuged for debris removal. **(B)** MES-CM sterilization and addition. The media is sterilized with a 0.2 µm filter and combined with cDMEM (1:1-1:7 dilution). Media is added to a hepatic spheroid and incubated with media changes every three days.

7.3. Dilute the fibroblast-conditioned growth medium with complete growth medium at a 1:1 to 1:7 dilution ratio and add to the desired experiment (**Figure 4B**).

NOTE: Utilize HFF/MRC-5 conditioned-media in the droplet formation assay.

### 8. HepG2 and HFF/MRC-5 Mixed Spheroid Formation

NOTE: Refer to steps 3.1 – 3.5. for HepG2 cell suspension preparation and steps 4.1. – 4.5. for HFF/MRC-5 cell suspension preparation.

8.1. Suspend the HepG2 cells in fresh cDMEM to achieve a final concentration of 1.0 x 105 cells/mL for each cell suspension.

8.2. Suspend the HFF/MRC-5 cells in fresh cDMEM to achieve a final concentration of 1.0 x 105 cells/mL for each cell suspension.

8.3. Transfer HepG2 and HFF/MRC-5 cell suspensions to a 15 mL sterile conical centrifuge tube at a 1:1 ratio to obtain a final concentration of 2.0 x 10^5^ cells/mL. Thoroughly mix the cell suspension and transfer 100 µL of cell suspension per well to the agarose-coated 96-well plate.

NOTE: The cell suspension density is set to obtain 20,000 cells per well in the agarose-coated 96-well-plate.

8.4. Centrifuge the plate at 340 *x g* for 10 minutes and incubate at 37℃ and 5% CO2 for 1-2 days (**Figure 5C**).

**Figure 5.**
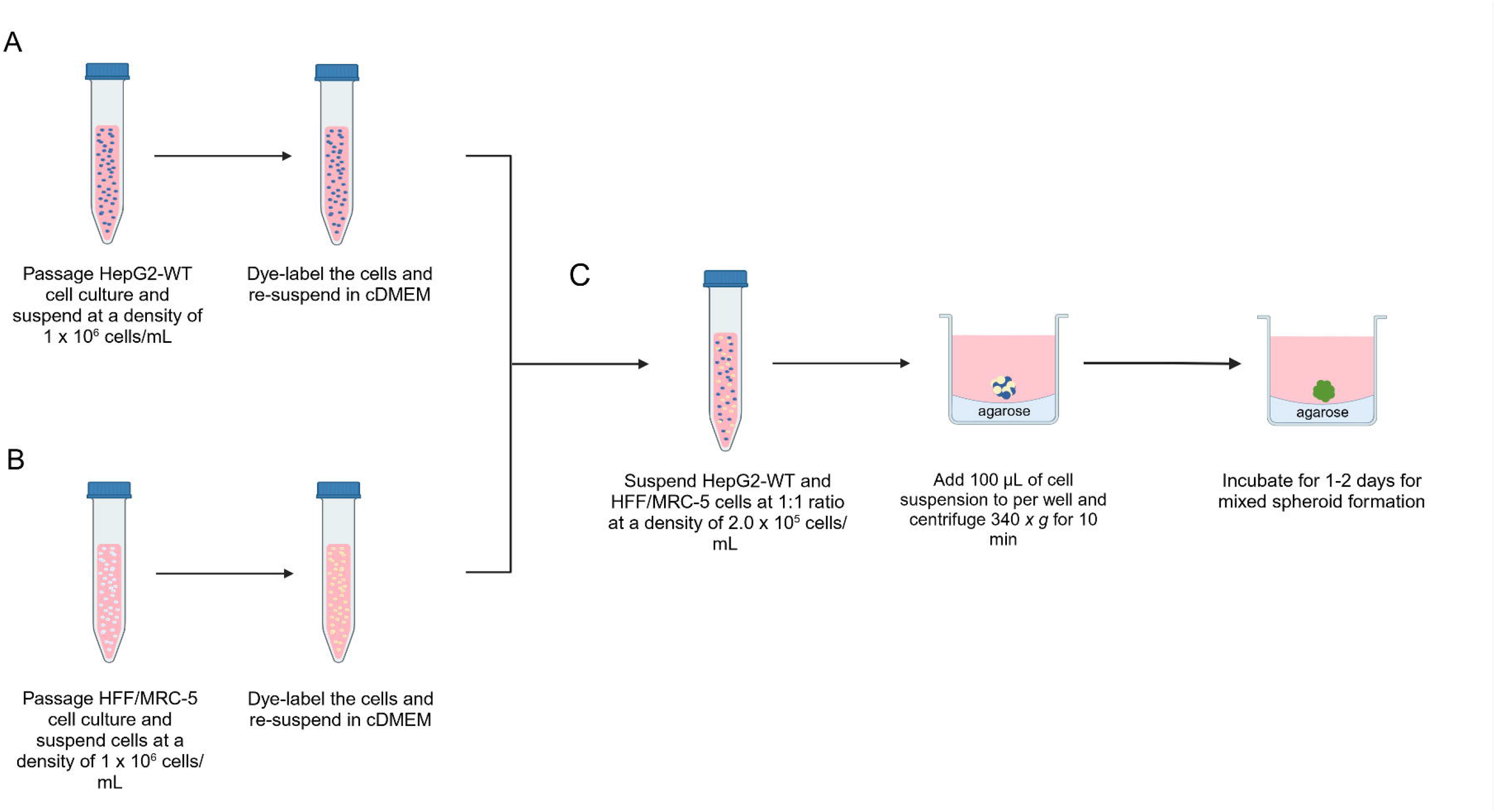
Schematic of Hepatic and Mesodermal-derived Mixed Spheroid Formation Assay. **(A)** Preparation of hepatic cell suspension. HepG2 cells are harvested and dye-labeled. **(B)** Preparation of Mesodermal-derived cell suspension. HFF/MRC-5 cells are harvested and dye-labeled. **(C)** HEP-MES mixed spheroid formation. HepG2 and HFF/MRC-5 cells are mixed at a 1:1 ratio at a final concentration of 2.0 x 10^5^ cells/mL and plated. The plate is incubated for 1-2 days until mixed spheroid formation is complete.

8.5. Addition of Fibrin Hydrogels

NOTE: This is an optional step that is done in 72 hours after plating HEP-MES mixed spheroids.

8.5.1. In a 15 mL sterile conical centrifuge tube, mix fibrinogen (3.25 mg/mL) and thrombin (12.5 units/mL) at a 4:1 ratio for the formation of fibrin and put the tube on ice.

8.6. Transfer the fibrin to each well with a pre-existing mixed spheroid at a 1:1 dilution with current cDMEM in the well. Incubate the plate at 37℃ and 5% CO_2_ for 30 minutes.

8.7. After incubation, add 75 µL of fresh cDMEM to each well and incubate at 37°C at 5% CO_2_ for 1-2 days.

NOTE: Change medium every other day after plating, gently removing 50% of cDMEM and replacing it.

### 9. High Density HFF/MRC-5 Migratory Model

NOTE: Refer to section 3 for HepG2 spheroid preparation **(Figure 6A**) and steps 4.1. – 4.5. for HFF/MRC-5 cell suspension preparation (**Figure 6B**).

**Figure 6.**
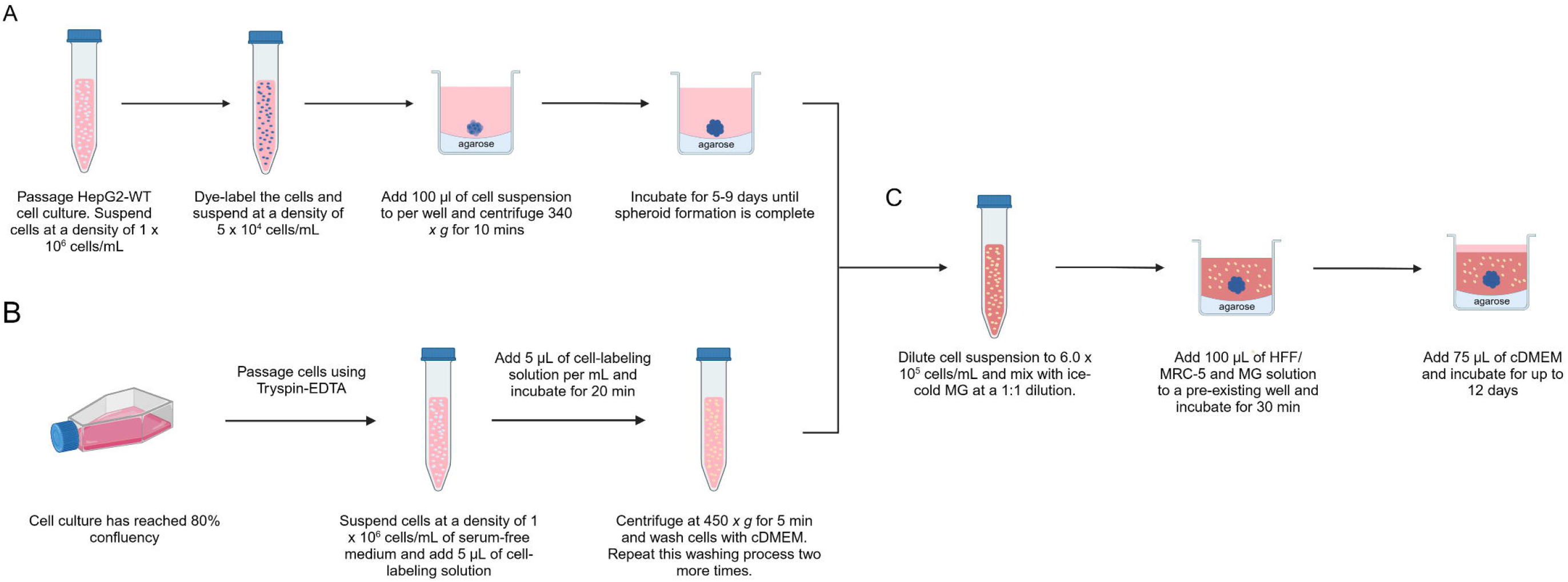
Schematic of High Density HFF/MRC-5 Migratory Model. **(A)** Hepatic spheroid formation. HepG2 cells are dye-labeled and plated for spheroid formation. **(B)** Preparation of Mesodermal-derived cell suspension. HFF/MRC-5 cells are harvested and dye-labeled. **(C)** HFF/MRC-5 cell suspension is pre-mixed with MG solution and added to pre-existing HepG2 well. The plate is incubated for 30 minutes and 75 µL of cDMEM is added. The plate is incubated for up to 12 days.

9.1. Suspend the HFF/MRC-5 cells in fresh cDMEM to achieve a final concentration of 6.0 x 10^5^ cells/mL and put on ice.

9.2. Mix the HFF/MRC-5 cell suspension and ice-cold MG/CG at a 1:1 dilution (**Figure 6C**).

NOTE: The cell suspension density is set to obtain 30,000 cells per well in the agarose-coated 96-well-plate.

NOTE: Thaw MG overnight on ice in a 2°C to 8°C fridge. Ensure pre-cooled pipettes, tips, and tubes when handling MG, as it forms a gel above 10℃.

9.3. Transfer 100 µL of HFF/MRC-5 and MG solution using a pipette to a well containing a HepG2 spheroid. Incubate the plate at 37℃ and 5% CO_2_ for 30 minutes.

9.4. After incubation, add 75 µL of fresh cDMEM to each well and incubate at 37°C at 5% CO_2_ for up to 12 days.

## REPRESENTATIVE RESULTS

Currently, increased interest exists in synthetic tissues for various biomedical applications, including disease modeling, drug discovery, and tissue engineering (**Figure 7**). In this field, human pluripotent stem cell (hPSC)-derived organoids, spheroids, and cells can be transformed into more synthetic, complex, and larger tissues. Achieving this involves applying principles of morphogenesis, utilizing tools like microfabrication, and incorporating biomatrices to engineer synthetic tissues more precisely (**Figure 7**). Several methods are presented here with this theme in mind.

**Figure 7.**
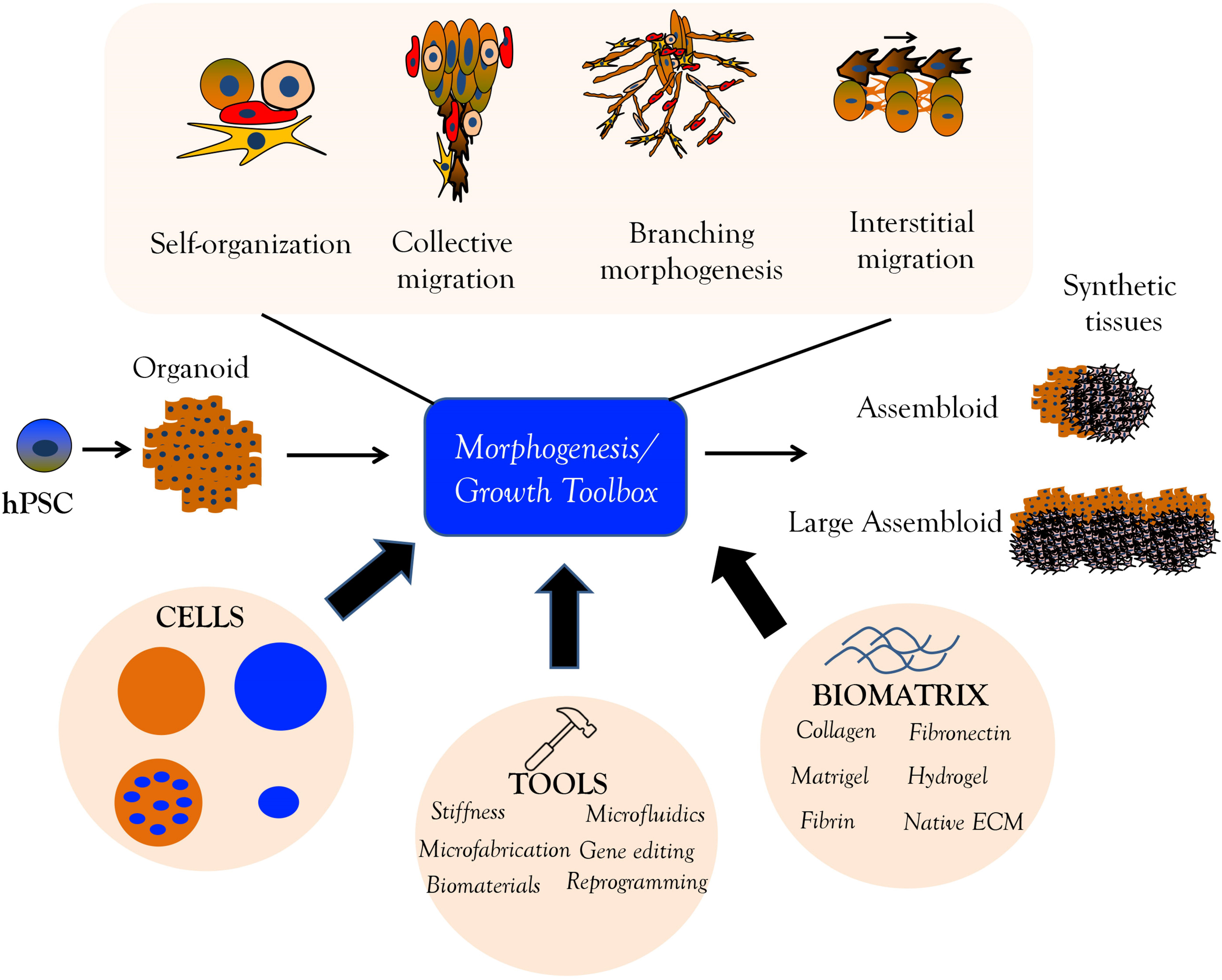
Overview. Based in the science of morphogenesis (top), stem cells will be implemented to build assembloids. The toolbox (middle), through the use of cells, tools, and biomatrix, and together with aspects of morphogenesis, can together be used to generate new synthetic tissues and assembloids.

### Effects of clustering on spheroid formation

The methods developed for this study were contingent upon successful 3D spheroid formation. Successful spheroid formation is indicated by cells fusing within five to nine days, forming a full spheroid. An early indication of success is cell clustering in the center of the well post-centrifugation of the cultivation plate. While spheroid formation occurred less frequently when cells were initially scattered, it still demonstrated that successful spheroid formation could occur. Unsuccessful spheroid formation is characterized by cells not spreading and fusing within nine days post-plating.

Spheroids of two different sizes were used to perform this experiment. Small spheroids (S) were plated at a concentration of 1,500 cells per well, and large spheroids (L) were plated at a concentration of 3,000 cells per well. HEP spheroids compacted and fully cultured by day five, irrespective of the spheroid size. The cell density per well did not affect the rate at which the spheroid formed but had a significant impact on spheroid size (**Figure 8A**). In this instance, it is crucial to observe an increase in opacity, indicating thickening of the initial disc-shaped tissue, which transitions from being more translucent to a spheroid configuration with increased opacity. HEP spheroids are uniformly less round and increase in optical density (increasing opacity) over time. If these HEP spheroids are still translucent, the spheroids should be either discarded or the experiment should be re-done. The same experiment was performed for HFF/MRC-5 spheroids, in which, the mesenchymal-derived spheroids compacted and fully cultured within 24 hours, irrespective of spheroid size. The cell density per well did not affect the rate at which the spheroid had formed (**Figure 8B**). Unlike hepatic spheroids, mesodermal-derived spheroids compact within 24 hours, irrespective of cell seeding density. Due to their contractile properties, MES spheroids are uniformly round and optically dense spheroids. Notably, HFF/MRC-5 cells compact much tighter than HepG2 cells, likely resulting in little impact of cell density on spheroid size (**Figure 8C**).

**Figure 8.**
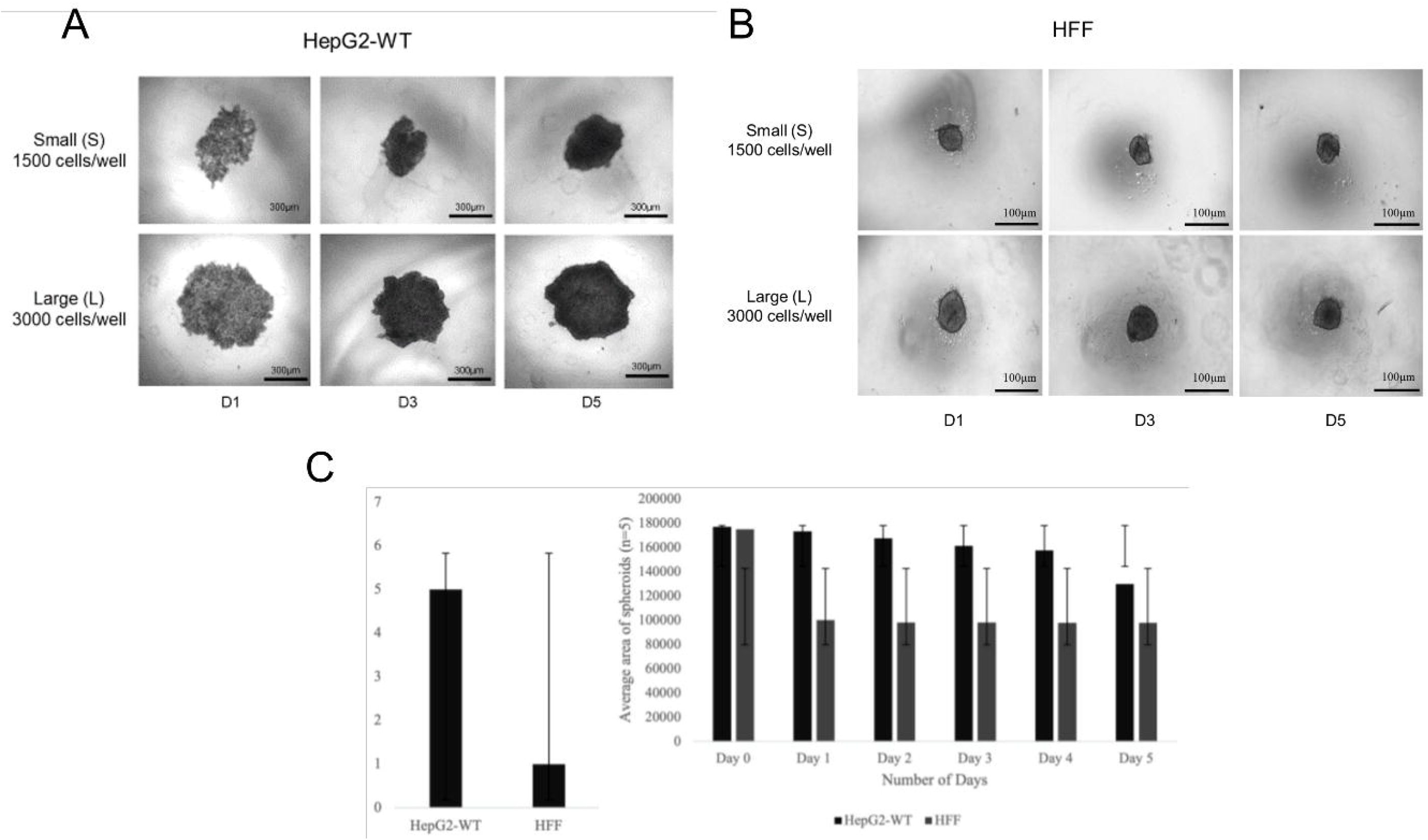
Compaction time of varying spheroid size. **(A)** Phase-contrast images of small (1,500 cells) and large (3,000 cells) hepatic spheroids on days 1, 3, and 5. Bar = 300 µm. **(B)** Same study as **(A)** except HFF cells. Bar = 100 µm. **(C)** Bar graph comparing HepG2 and HFF spheroid compaction. (n = 5). Right: bar graph plotting area of HepG2 and HFF spheroids over time. (n = 5).

### Factors that affect hepatic and mesenchymal (mesodermal-derived) assembloid formation

Hepatic and mesodermal-derived spheroids of varying sizes underwent cultivation to investigate the impact of size on the compaction time of spheroid formation. Hepatic and mesodermal-derived spheroids were co-cultured in GFR Matrigel (MG) or Collagen (CG) gel at a 1:5 dilution with complete growth medium (cDMEM) or fibroblast-conditioned medium (MES-CM) to explore the potential influence of matrix and conditioned medium on assembloid formation. Both cell types were dye-labeled before spheroid formation to illustrate the interaction between spheroids. In all experiments in which spheroids were dye-labeled, they were visualized using standard filters for green (488 nm), blue (450 nm), and red (594 nm). Images of the cells were captured using a Zeiss Axio fluorescent microscope equipped with Axiovision software to zero the background noise and images were analyzed using Image J. Phase-contrast, individual, and overlay images were obtained as .zip files, which were formatted into .tiff files using Image J.

It was observed that assembloid formation occurs regardless of matrix and medium (**Figure 9**). The effects of CG, MG, and MES-CM were studied, and assembloid formation was observed in all cases, although morphological details varied slightly (**Figure 9**). Details regarding these images will be re-used in later figures. To assess the impact of inter-spheroid distance on assembloid formation, distance measurements were taken along with the success of assembloid formation in the same experimental system (MG and DMEM). It was noted that the compaction time of an assembloid is directly proportional to the initial distance between hepatic and mesodermal-derived spheroids (**Table 1**).

**Figure 9.**
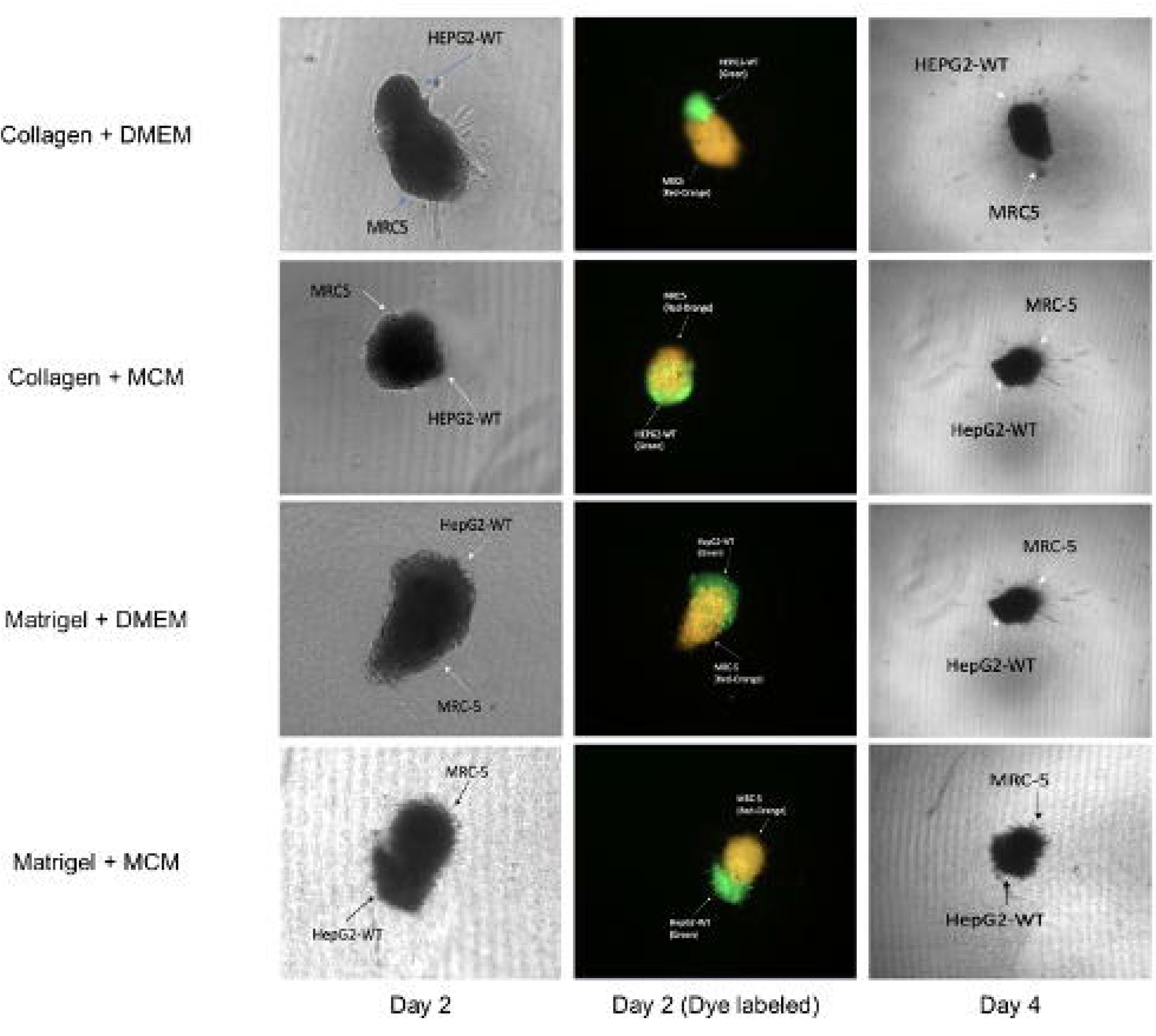
The effect of media and matrix on assembloid formation. Phase contrast and fluorescent images of fused HepG2/MRC-5 spheroids exposed to varying matrices (1:5 dilution) and growth medium: DMEM in CG, MES-CM in CG, DMEM in MG, and MES-CM in MG. Left: phase-contrast (day 2) images of fused HepG2/MRC-5 assembloid. Bar = 100 µm. Middle: fluorescent (day 2) of fused HepG2/MRC-5 assembloid. Bar = 200 µm. Left: phase-contrast of fused HepG2/MRC-5 assembloid. Bar = 200 µm. (green: HepG2; orange: MRC-5; arrows indicating spheroid types).

**Table 1.** The effect of distance on assembloid formation. Hepatic spheroids and mesodermal- derived spheroids were transferred to an agarose-coated well. The cells were dye-labeled prior to spheroid formation to demonstrate the interaction between spheroids. The spheroids were suspended in a 1:5 solution of the MG matrix suspended in cDMEM. The initial distance between the spheroids was measured using ImageJ and the corresponding compaction time of the assembloid formation was observed. The compaction time of assembloid formation is directly proportional to the initial distance between the spheroids. (n = 3).

| Growth Medium & Matrix | Initial Distances (μm) | Days to Merge |
| --- | --- | --- |
| Matrigel + DMEM | 492.742 | 6 |
| Matrigel + DMEM | 580.142 | 8 |
| Matrigel + DMEM | 974.051 | 12 |

### Building more complex assembloids with arm-like structures

Methods were developed to construct assembloids with branching cords (**Figure 10A**). To perform this experiment, hepatic spheroids were combined with biomatrix (GFR MG) in 384- well plate and surrounded by single fibroblasts at high density. These fibroblasts clustered, providing guides toward which hepatic cells migrated, thickening over time and forming cords. Hepatic spheroids were cultured in the MG droplet system containing a high density of MRC-5 cells (30,000 cells) in 100 µL of cDMEM and matrix in 384-wells, displaying small clusters of MRC-5 cells in the MG (**Figure 10B, days 3-4**). Subsequently, liver spheroids formed thick migrating strands protruding towards the fibroblast clusters, creating thick strands containing both cell lines (**Figure 10B, days 9-12**). This approach resulted in longer arms or cords, likely containing a mix of hepatic and fibroblast cells. Another approach involved constructing bridges or small interconnections (arms) between spheroids. To build small armed-structures, a larger HEP spheroid could be co-cultured in 384-well plate with a smaller mixed spheroid (**Figure 10C**), leading to small, knob-like arms. In summary, two approaches for forming additional arms to spheroids are presented.

**Figure 10.**
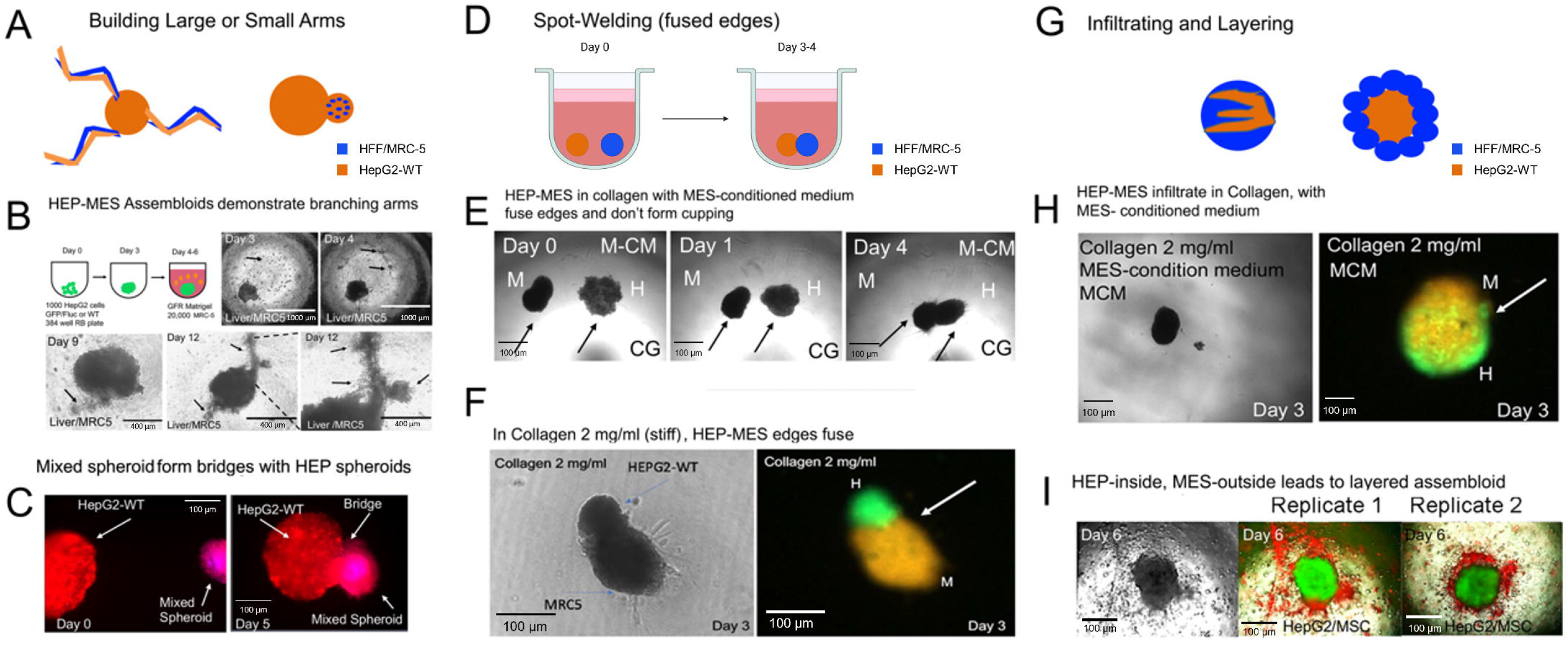
Engineering assembloids arms, junctions, and layers. **(A)** Schematic of building of large and small arms onto hepatic spheroids. **(B)** Phase-contrast images of HepG2 spheroids suspended in the MG droplet system, bearing high density (30,000 cells) of MRC-5 cells. Top: days 3, 4, 9, and 12. Day 3: MRC-5 cells initially after seeding (arrow). Bar = 1,000 μm. Day 4: MRC-5 cells spreading and interconnecting (arrows). Bar = 1,000 μm. Day 9: Thick hepatic cord (arrows). Bar = 400 μm. Day 12: multiple, thick hepatic cords. Bar = 400 μm. Bottom: images of a separate experimental replicate on day 12 demonstrating thick hepatic cord formation (arrows). Bar = 400 μm. Subfigures obtained with permission from Frontiers Journal^17^. **(C)** Fluorescent images of hepatic spheroid co-cultured with a HEP-MES mixed spheroid on days 0 and 5. Day 0: HepG2 and mixed spheroid initially after plating. Bar = 100 µm. Day 5: mixed spheroid has fused to the edge of HepG2 spheroid. Bar = 100 µm. (need to add dye-labeling information) **(D)** Schematic of building HEP-MES assembloids with fused edges (spot-welding). (arrows indicating spheroid type). **(E)** Phase-contrast images of HEP and MES spheroids co- cultivated in the CG droplet system exposed to MES-CM. Day 0: HEP and MES spheroids initially after plating. Bar = 100 µm. Day 1: HEP and MES migrating towards one another, but no fusion has occurred. Bar = 100 µm. Day 4: HEP and MES spheroids have fused at the edges of the spheroids. Bar = 100 µm. **(F)** Phase-contrast and fluorescent images of HEP-MES assembloid cultivated in CG droplet system (2 mg/mL) on day 3 demonstrating fused edges. (green: HepG2; orange: MRC-5; arrow indicating where fusion occurred). Bar = 100 µm. **(G)** Schematic of building HEP-MES assembloids with infiltration or a surface-MES layering. **(H)** Phase-contrast and fluorescent images of HEP-MES assembloid cultivated in CG droplet system (2 mg/mL) suspended in MES-CM on day 3 demonstrating infiltration of spheroids. (green: HepG2; orange: MRC-5; arrow indicating where fusion occurred). Bar = 100 µm. **(I)** Phase-contrast and fluorescent images on day 6 of HepG2 spheroid and MSC in MG droplet system. The right columns on the right are replicates 1 and 2 with fluorescent images on day 6. (green: HepG2; red: MSC). Bar = 100 μm. Subfigures obtained with permission from Frontiers Journal^17^.

### Spot-welding (fused edges) of complex assembloids

Methods were employed to construct assembloids with fused edges. The methodology to perform these experiments was HEP-MES assembloid formation assay, however, the MES spheroid had a lower density than previous experiments. Hepatic, and mesodermal-derived spheroids were positioned in a rigid CG environment (2 mg/mL) **(Figure 10D)**. In this rigid setting, instead of cupping, spheroids were observed to fuse at the edges, creating assembloids with indications of short arms or spot-welding (**Figure 10E**). On the other hand, described below, a cupping mechanism was also observed using the same culture system, in which the HEP spheroid surrounds the MES spheroid to form a partially fused assembloid. Regarding the spot-welding mechanism, similar data were observed with MRC-5 fibroblasts in CG (**Figure 10F**).

### Infiltrating and layering of complex assembloids

During liver organogenesis, in the developing liver diverticulum, HEP cells are surrounded by mesenchyme. Ultimately, they migrate or infiltrate into the mesenchyme (**Figure 10G**). In MES-CM and CG conditions, the addition of MES and HEP spheroids results in a different type of fusion, where an infiltrative pattern is observed (**Figure 10H**). Furthermore, a layering pattern can be achieved by placing single human mesenchymal stem cells (hMSC) in GFR MG at high density. They migrate towards a HEP spheroid and layer on the surface without infiltration, as shown with dye-labeling of the hMSC (**Figure 10I**). It is noteworthy that this is a different phenotype than when HFF was employed. The latter arrangement is observed in the liver diverticulum stage and many other endoderm-derived tissues. Two approaches are presented here, relevant for modeling the developing liver diverticulum and creating aspects of synthetic tissues.

### Fused hepatic and mesenchymal (mesodermal-derived) assembloid formation

The fusion of two spheroids to form assembloids is crucial for modeling the liver bud. This is because when two spheroids fuse, the cells likely migrate on top or between other cells, a phenomenon referred to as interstitial migration. Interstitial migration occurs in the liver bud when early migrating hepatoblasts move through the mesenchyme, potentially on top of other cells. Therefore, the fusion of two spheroids serves as a model for interstitial migration during liver organogenesis. Methods were developed to construct assembloids that fuse completely with separate layers (**Figure 11A**). HEP and MES (MRC-5) spheroids in MG form a fused assembloid by day 9 (**Figure 11B**). Dye-labeling analysis demonstrated that MES tissue remained inside, while the HEP tissue remained outside (**Figure 11C**). Importantly, the final spheroid is approximately the same size as the original spheroids, suggesting that the cells are packed at high density. Notably, the same phenomena occur in the absence of MG in 384-well plates when multiple MES spheroids are placed with a single HEP spheroid (**Figure 11D**). HEP- MES assembloids, in the absence of matrix, also form under low serum (2% FBS) and extremely low serum (0.2% FBS) conditions (**Figure 11E**). We then determined that at large distances (approximately 3 diameters of the spheroid), assembloids did not form (**Figure 11F**). To determine how spheroid composition determines spheroid fusion, it was demonstrated that mixed spheroids (containing MES and HEP cells) can fuse with MES spheroids by day 5, accompanied by an increased packing density, as expected (**Figure 11G**).

**Figure 11.**
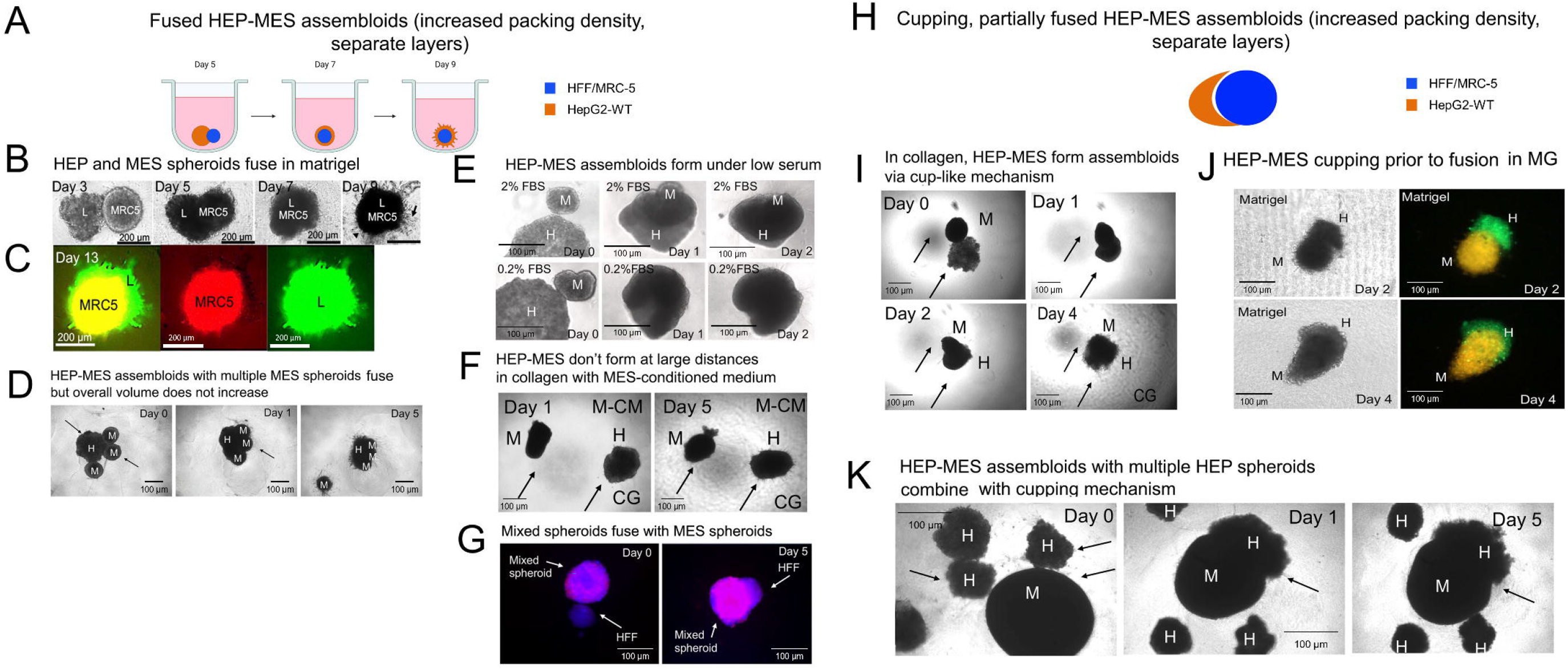
Partially and completely fused HEP-MES assembloids. **(A)** Schematic of building fused HEP-MES assembloids with increased packing density consisting of separate layers. **(B)** Phase-contrast images of HepG2 and MRC-5 spheroids co-cultivated in MG droplet system on days 3, 5, 7, 9. Days 3-5: HepG2 and MRC-5 spheroids interact. Day 7: migration at the spheroids edge and fusion occurred. Day 9: spheroid fusion and assembloid formation; arrows demonstrate protrusions. Bar = 200 µm. Subfigures obtained with permission from Frontiers Journal^17^. **(C)** Fluorescent images of HEP-MES assembloid, the same study as **(B)** on day 13. Day 9: spheroid fusion and assembloid formation; arrows demonstrate protrusions. (green: HepG2; red: MRC-5). Bar = 200 µm. Subfigures obtained with permission from Frontiers Journal^17^. **(D)** Phase-contrast images of HepG2 spheroid co-cultured with multiple mesodermal-derived spheroids on days 0, 1, and 5. Day 5: HEP-MES assembloid formation occurred but overall volume did not increase. **(E)** Phase-contrast images of HepG2 and MRC-5 spheroids co- cultivated in MG under low serum conditions (0.2% and 2% FBS) on days 0-2. Top: HEP-MES assembloid fusion occurs in 2% FBS by day 2. Bar = 100 µm. Bottom: HEP-MES assembloid fusion occurs in 0.2% FBS by day 2. Bar = 100 µm. **(F)** Phase-contrast images of HepG2 and MRC-5 spheroids co-cultured in the CG droplet system exposed to MES-CM. Day 0: HEP and MES spheroids initially after plating. Bar = 100 µm. Day 5: HEP and MES spheroids have not fused. Bar = 100 µm. **(G)** Fluorescent images of MES spheroid co-cultured with a HEP-MES mixed spheroid on days 0 and 5. Day 0: HFF and HEP-MES mixed spheroid initially after plating. Bar = 100 µm. Day 5: HFF and HEP-MES mixed spheroid fuse. Bar = 100 µm. **(H)** Schematic of building partially fused HEP-MES assembloid with cupping. **(I)** Phase-contrast images of HEP and MES spheroids co-cultured in the CG droplet system on days 0, 1, 2, and 4. Day 0: HEP and MES spheroids initially after plating. Days 1-2: HEP and MES spheroids interact. Day 4: HEP-MES assembloid formation occurs in a cup-like mechanism. Arrow indicating spheroid position during fusion. Bar = 100 µm. **(J)** Phase-contrast and fluorescent images of HEP-MES assembloid cultivated in MG droplet system on days 2 and 4. Top: HEP and MES spheroid interaction occurs. Bottom: partial fusion of HEP-MES assembloid. (green: HepG2; orange: MRC-5; arrow indicating where fusion occurred). Bar = 100 µm. **(K)** Phase-contrast images of MES spheroid co- cultured with multiple HEP spheroids on days 0, 1, and 5. Day 0: MES and HepG2 spheroids initially after platting. Day 1: HEP and MES spheroid interaction occurs. 5: HEP spheroid forms a cup around MES spheroid.

### Partially fused or cupping in complex assembloid formation

Successful assembloid formation in MG demonstrates several phenotypes, including the observation that the HEP spheroid undergoes “cupping” of the MES spheroid to form a partially fused assembloid (**Figure 11H**). This also establishes a visual model of interstitial migration and points toward the current mechanism of fused assembloid formation. Time-series studies of HEP and MES spheroids demonstrate that in CG, fusion occurs via a cup-like mechanism (**Figure 11I**). This is further demonstrated using dye-labeling (**Figure 11J**). Finally, the same phenomena are clearly illustrated when multiple HEP spheroids are used with a large, single MES spheroid (**Figure 11K**). The significantly smaller HEP spheroid cannot completely encompass the larger MES spheroid; therefore, it results in the same cupping mechanism. The data suggest that cupping without fusion can occur in MG or in cases where the MES spheroid is much larger than the HEP spheroids.

### Induced hepatic branching and linear migration via mesenchymal (mesodermal-derived) conditioned growth medium

Collective migration is a pivotal morphogenetic process during liver organogenesis. Tools for inducing collective migration *in vitro* are described here. Successful droplet formation of HEP spheroids and the use of MES-CM demonstrate outgrowth branching from the HEP spheroids (**Figure 12A-B**). HEP spheroids cultivated in the MG droplet system show that MES- CM induces collective migration, and the data demonstrate a concentration-dependent effect (**Figure 12B, right**). Cellular strands protruding from the hepatic spheroid are present with small branching, thick strands, and multiple levels of branching (**Figure 12B, left**). On day 11, the protrusions increased with interconnections and sheet formation (**Figure 12B, right**). MES-CM is potent, as a 1:7 dilution or MES-CM present for one day still leads to migration (**Figure 12B, right**). It was hypothesized that the MES-CM induced cell migration via TGFβ signaling pathway. A83-01, a TGFβ pathway inhibitor, was incorporated into the MES-CM at varying concentrations, and migration was significantly inhibited in a dose-dependent manner (**Figure 12C**). To determine the effects of extracellular matrix on collective migration, HEP spheroids cultivated in the CG droplet system demonstrate that MES-CM induces cell migration. However, the protrusions were thin linear strands and less branching by day 7 compared to the migration observed in MG (**Figure 12D**). Fibrin hydrogels were also tested, and thin, hair-like, narrow, radial protrusions of both HEP and MES cells were observed (**Figure 12E**).

**Figure 12.**
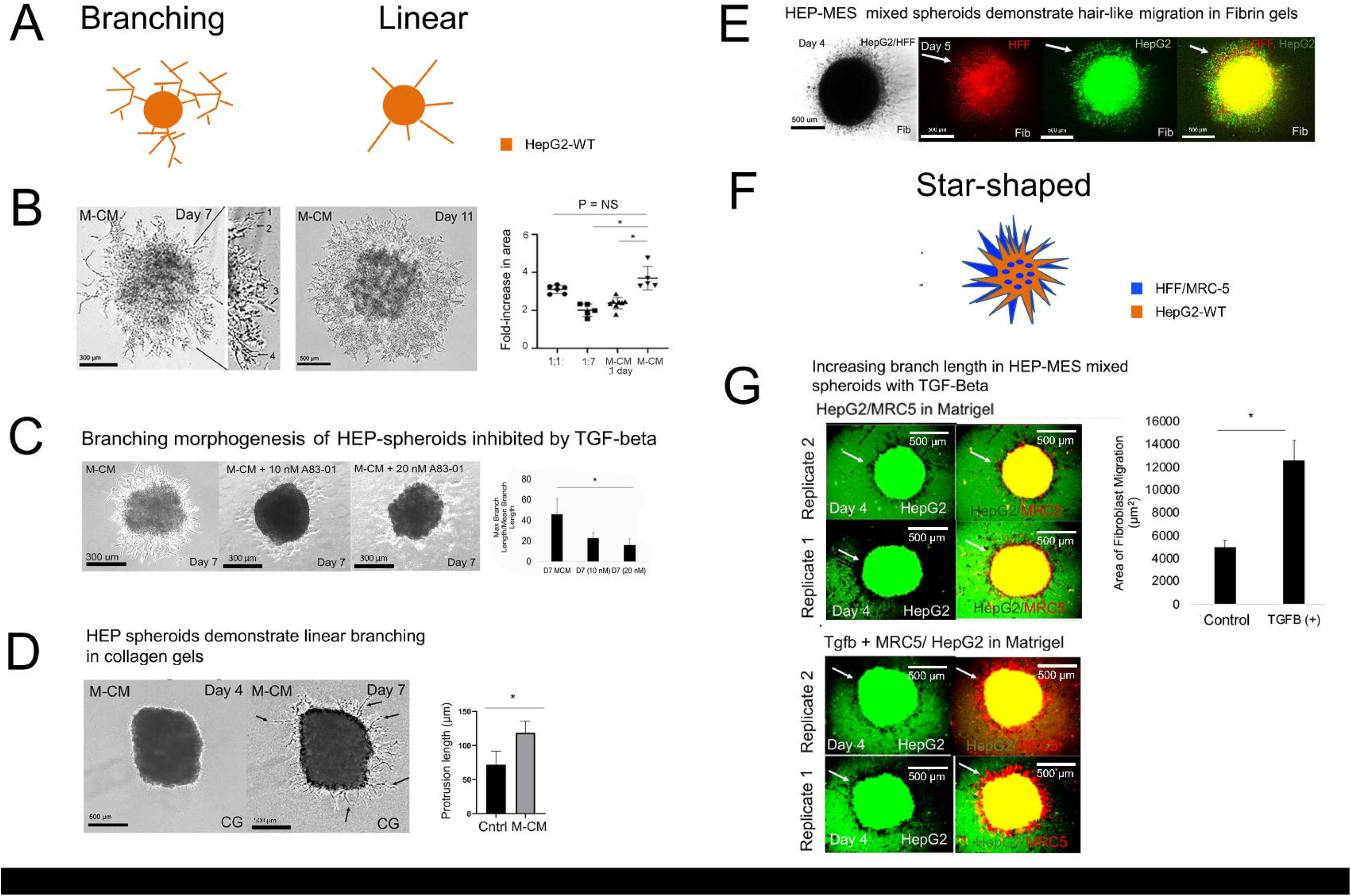
Methods for modulating collective migration from spheroids. **(A)** Schematic of branching and linear migration in HEP spheroids in the presence of MES-CM. **(B)** Phase-contrast images of HEP spheroid in the MG droplet system cultivated in varying dilutions of MES-CM on day 7 (low and high magnification) and 11. Day 7: HEP spheroid demonstrates various modes of 3D collective migration. Bar = 300 µm. Day 11: HEP spheroid outgrowth demonstrates interaction between branches. Bar = 500 µm. Plot comparing fold-increase in area across different dilution ratios of MES-CM (1:1, 1:7, MES-CM 1 day, and MES-CM).Comparison of MES-CM with 1:1 condition (P = 0.15, n = 3 for both conditions), MES-CM with 1:7 condition (P = 0.0056, n = 3 for both conditions), and MES-CM with MES-CM 1-day-only condition (P = 0.019, n = 3 for both conditions). Subfigures obtained with permission from Frontiers Journal^17^. **(C)** Phase-contrast images of HEP spheroid cultivated MG droplet system exposed to MES-CM alone, MES-CM with A83-01 (10 nM), and A83-01 (20 nM) on day 7. Bar = 300 µm. Bar graph analysis comparing day 7 MES-CM and MES-CM + A83-01 (20 nM) (P = 0.047, n = 3). Plotted means ± SD. Significance is defined as P ≤ 0.05. Subfigures obtained with permission from Frontiers Journal^17^. **(D)** Phase-contrast images of HEP spheroid cultured in the CG droplet system cultivated in MES-CM on days 4 and 7. Arrows specify thin filopodia-like extensions into the CG. Bar = 500 µm. Bar graph comparing protrusion length of HEP spheroids in cDMEM and MES-CM conditions (P = 0.012, n = 3). Subfigures obtained with permission from Frontiers Journal^17^. **(E)** Phase-contrast and fluorescent images of HEP-MES mixed spheroid cultivated in fibrin gels exposed to MES-CM on days 4 and 5. (green: HepG2; red: HFF; arrows indicated protrusions). Bar = 500 µm. Subfigures obtained with permission from Frontiers Journal^17^. **(F)** Schematic of star-shaped migration of HEP-MES mixed spheroids. **(G)** Fluorescent images of HEPG2-GFP (HepG2 cells expressing green fluorescent protein) and HEP-MES mixed spheroids co-cultivated in the MG droplet system after treatment with TGFβ1 (20 ng/ml) on day 4. Left: HepG2 cells in HEP-MES mixed spheroid exposed to TGFβ1 on day 4. Right: HEP-MES mixed spheroid exposed to TGFβ1 on day 4. Replicates 1 (below) and 2 (above) are shown. (green: HepG2; red: MRC-5; yellow: combined; arrows show HepG2 and MRC-5 migration). Bar = 500 µm. Bar graph comparing the area of fibroblast migration when exposed to TGFβ1 (HepG2- GFP/MRC-5 and HepG2-GFP/MRC-5 + TGFβ1, 20 ng/ml), P = 0.012, n = 3. Plotted means ± SD. Significance is defined as P ≤ 0.05. ∗ is used to denote the significance of experimental data. Subfigures obtained with permission from Frontiers Journal^17^.

### Inducing hepatic co-migration via MES conditioned medium

Mixed spheroids (HEP-MES spheroids) were utilized for collective migration, as a model of co-migration which occurs during early liver organogenesis (**Figure 12F**). These mixed spheroids in MG result in migration, and when TGFβ1 growth factor was added, it resulted in significantly increased collective migration (**Figure 12G**). Thus, co-migration can also be modeled with mixed spheroids.

## DISCUSSION

In this protocol, multiple methods are presented for cultivating both simple and complex assembloids, as well as techniques for inducing 3D collective cell migration in early liver organogenesis^14, 17^. Several critical steps are outlined in the presented protocols, with spheroid formation being a key aspect across all methods. Achieving spheroid formation involves using microwells (96- or 384-wells) with either non-adherent or agarose-coated plates. Spheroids are abundantly produced for subsequent experimentation, accounting for those that may not reach the expected size or have irregular shapes. Spheroids that do not meet the desired criteria are not utilized in downstream experiments. The initial efficiency depends on various factors, including the user. Considerable expertise is required to handle organoids, involving aspects such as spheroid (or organoid) formation, transfer between wells, addition of biomatrices upon spheroids, and placement of multiple spheroids per well.

Formation of the spheroids demands meticulous attention to repeatable cell counting and seeding, agarose coating (or non-adherent), regular medium changes, dye-labeling methods, and gentle handling of spheroids and plates, along with microscopy. Dye-labeling specifics, including cell number and type, as well as, the labeling duration, should be experimentally determined. Typically, concentration of MG averaged 9.02 ± 0.63 mg/mL. Furthermore, experience suggests that minor differences do not significantly impact phenotype, it could be highly beneficial to track protein concentrations for each MG lot in initial experiments. In terms of the CG, the concentration was prepared to be 2 mg/mL.

For instance, preparing the agarose solution necessitates strict adherence to maintaining appropriate concentrations, seeding wells with proper volumes to allow for the development of a meniscus (curvature), and enable cell collection in the center. Careful consideration of agarose concentration, volume, and gel solidification is crucial. Selecting the correct micropipette tip size and employing proper transferring techniques are essential for handling spheroids effectively. The use of glass wells can enhance visualization. As observed in the paper, spheroids initially appear as translucent discs, becoming more opaque over time, providing a means to monitor spheroid formation. Several factors can influence this process. Lastly, for assembloid formation^9^, precise transfer of spheroids is vital, placing them within a couple of spheroid diameters or less to effectively observe changes.

Here, a toolbox of methods has been established using cell lines to develop complex assembloids and migrating spheroids. It is noteworthy that cell lines were employed for method focus, but using primary cells or hPSC-derived cells would be advantageous for human personalized tissues, and our assembloid techniques have been validated with hPSC-derived HEP cells^14,15^. However, a significant limitation of these systems is the use of GFR MG, a mouse tumor extracellular matrix protein mixture primarily used in these cultivation models. However, it is a major limitation due to its tumor-derived origin and high cost. Additionally, the droplet formation system degrades significantly after two weeks, suggesting the implementation of alternative gels such as CG mixtures or sodium alginate. Furthermore, the current systems can only grow to the dimensions of the spheroids, but not beyond, emphasizing the need to identify and address limitations to growth.

The lack of methods to specifically study early events in early liver organogenesis^14^ and, more generally, methods to generate synthetic tissues^1^. These methods offer a toolbox for building assembloids and assembling larger, more complex tissues for enhanced *ex vivo* organ modeling. This holds potential for *in vivo* therapeutic approaches, in addition to modeling structures during organogenesis (liver bud) and applications in biopharma.

The protocols presented enable models of different modes of liver cell migration, including co-migration, interstitial migration, and branching morphogenesis, along with assembloids that collectively can be used to construct more complex tissues for various biomedical and biopharma applications.

## FIGURE PERMISSIONS

Permission to reproduce images from Frontiers Biotechnology and Bioengineering Journal were granted through the Creative Commons Attribution (“CC BY”) license 4.0.

## ABBREVIATION LIST

BMP4: Bone morphogenetic protein 4
FGF-2: Fibroblast growth factor 2
PROX1: Prospero homeobox 1
HBs: hepatoblasts
HEP: hepatic
HGF: hepatocyte nuclear factor 4 alpha
hMSC: human mesenchymal stem cells
HPSC: human pluripotent stem cells
MES: mesenchymal
MES-CM: fibroblast-conditioned medium
TBX3: T-box transcription box factor 3
TGFβ1: transforming growth factor beta1

## ACKNOWLEDGMENTS

Authors thank Jenna Venturo, undergraduate in the lab, for her assistance. NP was supported by the UB CBE startup funds, a Mark Diamond Fellowship, the New York State Stem Cell Science C024316, the UB the Stem cells in regenerative medicine (ScIRM) center, the University at Buffalo Center for Cell, Gene, Tissue Engineering (CGTE).

